# Three-State Gene Expression Model Parameterized for Single-Cell Multi-Omics Data

**DOI:** 10.1101/2025.07.16.665109

**Authors:** Thibaut Peyric, Thomas Lepoutre, Anton Crombach, Thomas Guyet

## Abstract

We present a novel three-state gene expression model designed to elucidate the underlying mechanisms of mRNA transcription and its regulation. Our model incorporates gene regulatory processes by explicitly including a transcription factor-bound state, thereby capturing the dynamic interplay between transcription activation and chromatin dynamics. We fit the model to paired single-cell ATAC-seq and single-cell RNA-seq data, as these data give us simultaneous information on a gene’s transcriptional state and its accompanying chromatin state. Working at the pseudo-bulk level, we extract biologically meaningful high-level descriptors from homogeneous cell (sub)populations, such as the mean and variance of gene expression as well as the fraction of accessible chromatin. Crucial to the computational feasibility of our approach, these descriptors can be analytically related to our model parameters. Despite the increased complexity needed to capture regulatory processes in our model, it remains sufficiently parsimonious to infer parameters reliably from experimental data. Each parameter has a clear biological interpretation, reflecting properties such as burst frequency, chromatin opening and closing dynamics, and basal or regulated expression. Fitting the model to a large collection of genes allows us to analyze the parameters and distinguish so-called gene expression strategies. The model parameters reveal a small number of distinct expression strategies among gene clusters, providing data-driven novel insight into context-dependent regulation of gene expression.

## 1 Introduction

Understanding gene expression is fundamental to molecular biology, as it governs cellular behavior, differentiation, and response to environmental stimuli. Gene expression refers to the process by which genetic information is transcribed from DNA into RNA and subsequently translated into proteins. Over the years, advancements in technology have revolutionized our ability to detect and quantify mRNA molecules, transitioning from bulk measurements to high-resolution single-cell analyses that capture the stochastic nature of transcription [4,19]. These innovations have shed light on the burst-like behavior of transcription, characterized by periods of rapid mRNA synthesis interspersed with quiescent phases [11]. To capture these dynamics, one approach is the two-state Markov model [1,17], which conceptualizes transcriptional activity as stochastic transitions between an active and an inactive state. In this framework, a gene switches between these two discrete states with specific transition probabilities, defining the temporal dynamics of mRNA synthesis. This probabilistic representation effectively encapsulates the inherent noise and burst-like characteristics of transcription.

However, despite the utility of this model, it lacks capabilities regarding inferring the regulatory mechanisms that influence gene expression. Gene regulation is a complex interplay of chromatin accessibility, epigenetic modifications, and transcription factors (TF). The absence of explicit regulatory state in the model limits its ability to explain gene expression variability under different cellular conditions. Understanding the TF and chromatin accessibility interactions would enable better integration of the diverse types of omics data currently available. While it is now possible to access to paired multi-omics dataset —such as transcriptomics, epigenomics, or chromatin accessibility— their full interpretation remains limited due to the complexity of their inter-dependencies [9].

To capture these mechanisms hiding behind gene expression, several studies have proposed generalized models with *N* states instead of only two. These generalized models offer greater flexibility to represent complex regulatory dynamics, such as multi-step activation processes and more accurate transcriptional bursts [7]. The two main approaches are either models where every state is capable of transcript expression [2,8,25], or models in which only a single state is [7,26]. Although these models offer valuable theoretical insights and capture a complex spectrum of gene expression dynamics, their inherent complexity and large number of parameters present significant challenges when fitting experimental data. Indeed, for an additional state introduced to a model of *N* states, between 2 and 2*N* new parameters are needed. On top of that, one must determine for each gene the appropriate number of states to accurately model it, which introduces an additional parameter to be determined. In short, the primary consideration is that the number of possible solutions fitting the data increases exponentially with the number of parameters in the model, significantly complicating the optimization process. That is why in many cases the required detailed parameterization hinders the direct application of these models to experimental single-cell data.

Finding ways to extract information from the many kinds of single-cell data that exist nowadays is crucial to better fit computational models and thus shed light on the regulatory mechanisms underlying gene expression. This is why the use of multi-omics datasets is particularly valuable, especially those combining single-cell RNA sequencing (scRNA-seq) and ATAC sequencing (scATAC-seq). Such datasets provide complementary information, capturing both gene expression levels and chromatin accessibility. This dual-layer view enables a more comprehensive understanding of gene regulation mechanisms and improves the identifiability and robustness of model parameters. However, the relationship between chromatin accessibility and gene expression remains poorly understood [5]. Integrating these two omics tries to disentangle their reciprocal influence, but at the same time highlights this entanglement. Indeed, while chromatin accessibility modulates transcriptional activity by controlling access to regulatory elements, gene expression itself can feedback to alter chromatin structure, highlighting a bidirectional and dynamic interplay between the two [23]. Thus RNA-seq and ATAC-seq offer complementary perspectives on cellular states, though inferring one from the other remains problematic [5]. Chromatin accessibility of promoters, as measured by ATAC-seq, does not consistently predict levels of gene expression, largely because it fails to capture the nuanced regulatory control that modulates transcription. Without accounting for regulatory control, both RNA and ATAC data fluctuate at too different a time scale for reliable interpretation [3,18].

In response to these challenges, we propose a three-state model that balances biological realism with practical applicability by integrating multi-omics datasets. Based on the well-known two-state model, our model incorporates one new regulation state. By limiting the number of states and parameters, our approach facilitates the integration of high-resolution single-cell RNA-seq and ATAC-seq data, enabling a more robust and accessible analysis of gene expression regulation at the single-cell level. Moreover, by combining features of transcriptional and chromatin dynamics, our model provides a framework for interpreting (paired) single-cell multi-omics data.

## 2 Methods

In this section, we introduce a three-state gene expression model that is well-suited to be fit to paired single-cell multi-omics data (Figure 1). We will begin by examining the model itself, the hypotheses underlying its construction, and the biological interpretation of its parameters. Next, in order to fit the model to data (on a gene-by-gene basis), we leverage analytical expressions derived in Appendix A, which give us high-level descriptors of the model: the mRNA expression mean *M* ^*e*^ and variance *V* ^*e*^, and the fraction of open chromatin *F* ^*c*^. Then, for a given parameterization of the model, its fit to data is captured by a loss function (Figure 1, black box), which compares the model s high-level descriptors to their counterparts extracted from single-cell data. The loss function is minimized via gradient descent, allowing the model to reproduce gene expression in a manner that closely reflects the observed data.

**Fig. 1.**
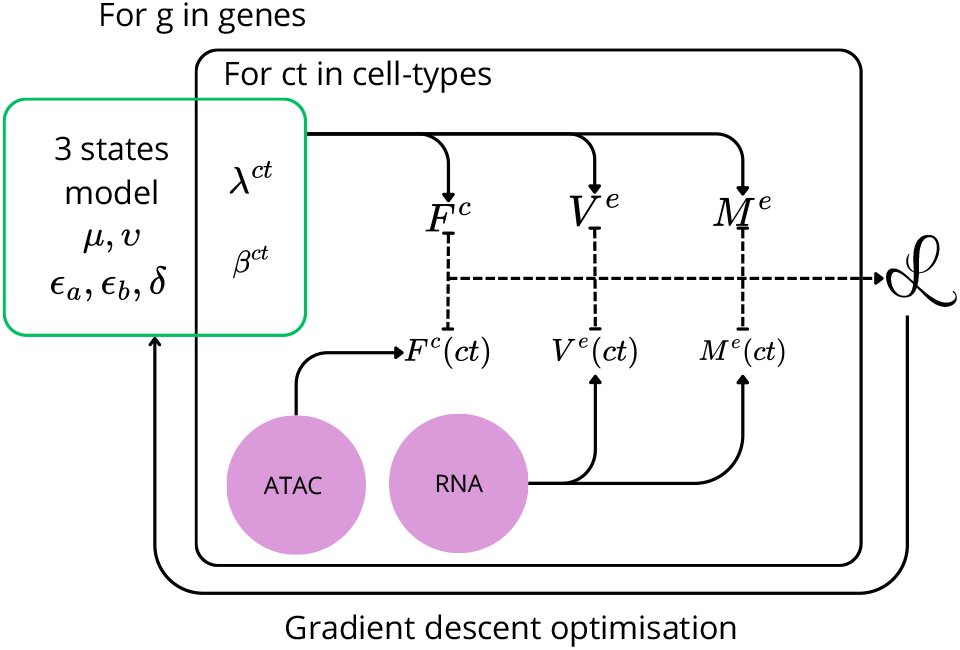
Schema of the three-state model fit to single-cell multi-omics data. For each gene, we define a three-state model with seven parameters (green box). Five parameters are fit on the basis of the whole population of cells, while the two remaining parameters, *λ*^*ct*^ and *β*^*ct*^, are fit per cell type (*ct*, black box). During fitting, performance of the model is encoded in a loss function, comparing three high-level model descriptors to their data-derived equivalents (purple circles), namely the fraction of open chromatin F, mean mRNA expression level M and variance of mRNA expression V. See main text for details.

### 2.1 A Three-State Model for Gene Expression

Our three-state model is designed with two goals in mind. Firstly, we aim to take into account recent hypotheses about gene expression mechanisms, and secondly, we want to limit the number of parameters to facilitate inference from data. We begin with a minimal two-state model, comprising an inactive state (*I*) and an active state (*A*), governed by the parameters *λ, µ, ϵ*_*a*_, and *δ*. Our first hypothesis posits that gene expression primarily varies through changes in burst frequency across different cell types [11]. This assumption leads to defining a distinct *λ* which represents the frequency of transcriptional bursts, for each cell type. While the other parameters are the same across all cell types (i.e., they are characteristic to a gene). Variations in *λ* are interpreted as differences in chromatin accessibility, influenced by structural factors such as nucleosome positioning and histone modifications. Under this hypothesis, gene expression is proportional to burst frequency and the model behaves as a linear regression [18].

The second hypothesis suggests that chromatin opening and closing is a much faster process than transcription, but that transcription factors can modulate this chromatin dynamic by stabilizing the open chromatin state [3]. To capture this, we introduce a new transcriptionally active state (*B*) that is disconnected from *I* and governed by transition probabilities *υ* and *β* (Figure 2). This addition allows the system to become “trapped” in a stably-open state *B*. We complete this part by introducing a cell-type specific *β* parameter, which captures context-dependent regulatory influences per cell type.

**Fig. 2.**
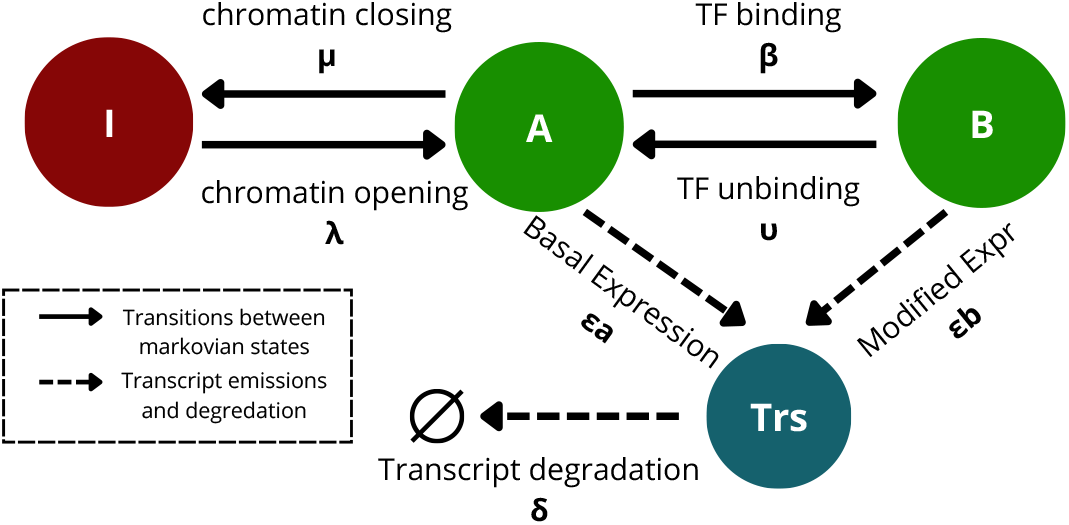
Representation of the three-state model for gene expression. There are three Markov states, one where a gene is inactive (*I*) and its chromatin closed (red circle); and two active states, *A* and *B* (green circles), where both correspond to open chromatin and *B* indicates a state of TFs influencing gene expression. Transition probabilities between the states are represented by *λ, β, µ*, and *υ*. Parameters *ϵ*_*a*_ and *ϵ*_*b*_ represent the probability to emit a transcript (*Tr*, blue circle) and to increase by one the total number of transcripts (*n*). *δ* is the probability for a single transcript to degrade, implying the total probability to decrease gene transcripts is *nδ*.

As a result, our Markov model is defined by a state space that is a pair (*S, n*) where *S* ∈ {*A, B, I*} is the gene state and *n* ∈ ℕ is the number of transcripts in the cell. The model transitions between an inactive gene state *I* with closed chromatin, an active gene state *A* with open chromatin, and another active state *B* with open chromatin bound by TFs. The model is governed by seven parameters including four for transitions between states (Figure 2). The parameter *µ* represents the probability of returning to an inactive state; the higher its value, the more the gene tends to be unexpressed. The parameter *υ* represents the probability of transitioning from a TF-bound state *B* to an unbound state *A* and reflects the affinity between a transcription factor and a gene. A higher *υ* value implies that a TF will spend less time in contact with the gene. The model emits transcripts (*Trs*) one by one with a probability of *ϵ*_*a*_ and *ϵ*_*b*_, corresponding to basal expression and regulated expression of the gene, respectively. Finally, *δ* defines the degradation probability of a transcript. In formula:

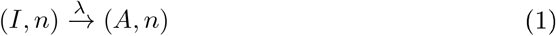

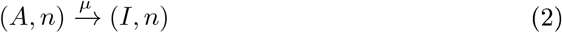

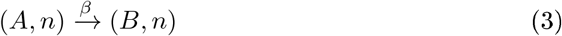

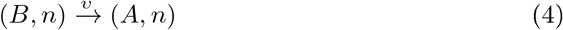

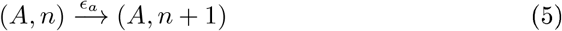

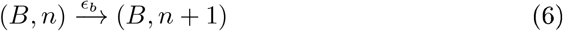

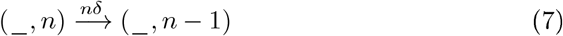

Of the seven parameters, five are considered intrinsic to each gene (*ϵ*_*a*_, *ϵ*_*b*_, *υ, µ* and *δ*) and we keep them the same across different cell types. These parameters encapsulate fundamental gene-specific properties such as basal transcription rates *ϵ*_*a*_, enhanced transcription rate *ϵ*_*b*_, TF binding affinities *υ*, chromatin remodeling dynamics *µ*, and degradation rate *δ*.

In contrast, parameters *λ* and *β* are assumed to be dependent on the cellular context. Each cell-type will have its own version if these parameters. *λ* represents the frequency of transcriptional bursts, which is closely linked to chromatin architecture and DNA compaction. Variations in *λ* across cell types reflect differences in chromatin accessibility, as determined by structural elements such as nucleosome positioning and histone modifications [6]. *β* encapsulates the effect on gene expression of external regulatory factors, such as transcription factors, signaling pathways, and epigenetic modifications. The parameter quantifies how a gene responds to its regulatory environment, potentially providing insight into differential gene activation across cell types.

### 2.2 Extraction of Descriptors from Single-Cell Multi-Omics Data

Single-cell RNA-seq and single-cell ATAC-seq data are notoriously difficult to integrate [5,10,16]. We propose to overcome this challenge by shifting perspective from the single cell to the pseudo-bulk level. By aggregating data across homogeneous populations of cells, we can obtain more stable and representative descriptors of gene expression and chromatin state, which are less sensitive to the asynchronous dynamics of chromatin and transcription inherent to the single-cell level. Indeed, chromatin is substantially more dynamic than gene expression, capable of transitioning between open and closed states within milliseconds to seconds [3]. In contrast, transcriptional bursts occur at much lower frequencies, ranging from minutes to several hours [11]. This discrepancy in timescales complicates a direct association between chromatin accessibility and transcription at the single-cell level. An individual cell may exhibit high transcript levels, while its chromatin is already closed, or vice versa [15]. Thus to overcome timescale complications, we want to describe genes at the level of a group of cells.

To this end, we extract three “high-level” statistical descriptors from the scRNA-seq and scATAC-seq data. First, we preprocess scRNA-seq data according to a standard Seurat-style approach [13] to obtain homogeneous clusters of cells [21]. By assuming homoscedasticity within such cell clusters, we can treat the collective behavior of these cells as representative of a single gene expression trajectory over time (Figure 3). Indeed, at this level gene expression follows a Negative Binomial distribution. This distribution is fully parameterized by its mean and variance, allowing scRNA-seq data to completely characterize the statistical properties of the distribution of gene expression within a population. In other words, as descriptors of transcriptional activity we compute for each gene the mean, *M* ^*e*^, and variance, *V* ^*e*^, of its mRNA expression level in each cell cluster.

**Fig. 3.**
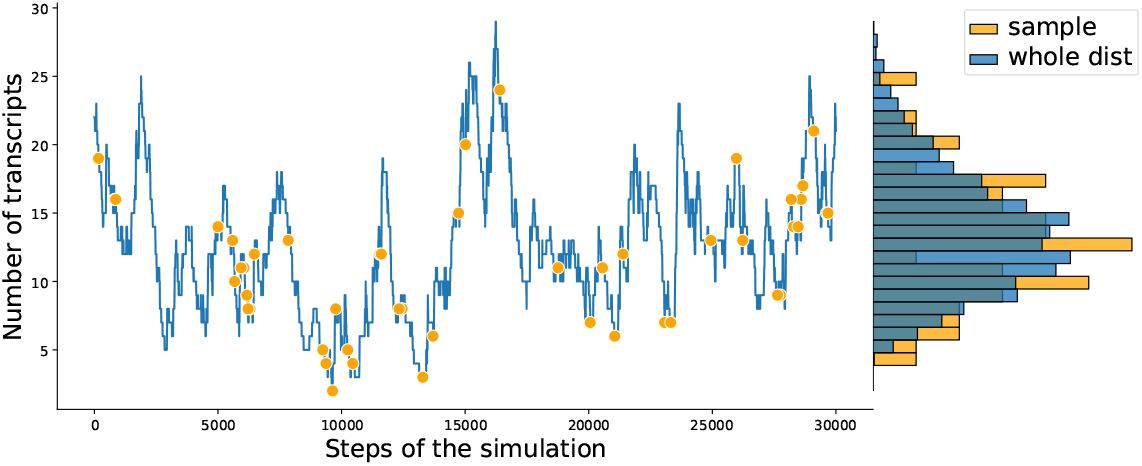
Simulation of gene expression by the three-state model. **Left** Under the homoscedasticity assumption that gene expression variance remains constant over time, we can use a population of homogeneous cells (orange dots) as representing snapshots of the temporal expression dynamics of a single cell (blue line). **Right** Histograms of the whole transcript distribution (blue) and a sampling (orange).

These two descriptors, however, do not capture the temporal, burst-like, nature of gene expression. To address this limitation, we use scATAC-seq data, which measures chromatin accessibility. Specifically, scATAC-seq data provide a list of accessible DNA regions for each individual cell. In our current analysis, we retain only the regions located at the Transcription Start Site (TSS) of each gene. This results in a binary cell-by-gene matrix indicating chromatin accessibility at gene promoters. Next, we use the cell clusters derived from the corresponding scRNA-seq data to compute the average chromatin accessibility for each gene within a cluster. This is our third high-level descriptor, an estimation of the mean fraction of time the chromatin remains in an open state, *F* ^*c*^. And this fraction directly corresponds to the average time the gene spends in the active state (*A* or *B*) in our proposed model.

We note that we rely on paired scRNA and scATAC data, where the two types of experimental data are derived from the same cell or nucleus (i.e., matching barcodes). Due to the limited resolution of scATAC-seq data for defining cell types [22], it is most effective to use paired data and transfer the cell cluster information from the RNA data type to the other. Alternatively, independently generated scRNA-seq and scATAC-seq datasets can be used, if both are annotated with consistent cell-type labels.

### 2.3 Model Fitting using High-level Descriptors

We can now fit the model using the three high-level descriptors that are extracted from single-cell data as described in the above section and from the model itself through the descriptors analytical solutions (see Appendix A). Each gene is represented by a unique and independent model, characterized by its own set of parameters. The fitting process is based on a gradient descent optimization, where at each iteration we compare the descriptors, *F* ^*c*^, *M* ^*e*^, and *V* ^*e*^ derived from the data with those computed by the model, using a dedicated loss function. The loss function is a central component for fitting the model to data, as it defines both the optimization objectives and the relative priorities among them. We elaborate on it below.

For each gene, let *k* denote the number of cell types, and let *ct*^*i*^ represent the single cells corresponding to a cell type *i* for *i* in [*k*]. For each cell-type, *F* ^*c*^, *M* ^*e*^ and *V* ^*e*^ can be evaluated from the data, which provides us the true-values of *F* ^*c*^(*ct*^*i*^), *M* ^*e*^(*ct*^*i*^) and *V* ^*e*^(*ct*^*i*^). And they can be efficiently computed as a function of the parameters of the model given the analytical solutions of *F* ^*c*^(*λ*^*i*^, *β*^*i*^, *P*), *M* ^*e*^(*λ*^*i*^, *β*^*i*^, P) and *V* ^*e*^(*λ*^*i*^, *β*^*i*^, P) (see Appendix A).

Then, parameter inference consists in finding the model parameters that minimizes the error made between the real and inferred data. More specifically, we optimize the following loss function:

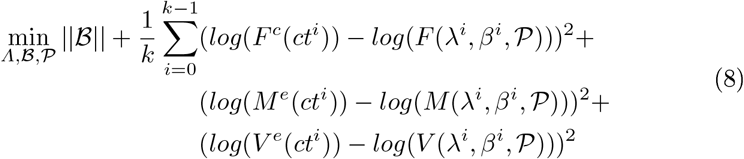

where *Λ* = {*λ*^0^, …, *λ*^*k−*1^} and ℬ = {*β*^0^, …, *β*^*k−*1^} are the parameters specific to each cell type and P = {*µ, υ, ϵ*_*a*_, *ϵ*_*b*_, *δ*} are the parameters shared across cell types.

We add an L1 regularization on *ℬ* which encourages the model to restrict gene regulation to a minimum. Thus, if a gene is not regulated by transcription factors, *β* will tend to 0. This means that our three-state model—designed to incorporate regulatory influences—defaults to the behavior of the two-state model, when regulatory effects are negligible. Moreover, by setting the regulatory parameter *β* to zero, the three-state model effectively reduces to the two-state one, ensuring that the linear relationship of gene expression being proportional to burst frequency is preserved. This model behavior is desirable for accurately capturing gene expression dynamics in cases where regulatory influences are minimal or absent. The logarithmic transformations in the three loss terms is motivated by a large range of magnitudes in gene expression values, which can span from 1*e*^*−*3^ to 1*e*^3^. In other words, the logarithmic transformation balances the error between all descriptors that have different scales.

In summary, the three-state model provides a mechanistic framework linking transcriptomic and chromatin accessibility features through three descriptors characterizing gene expression distributions. Although the model involves more parameters than directly observable features, incorporating multiple cell populations—each with shared parameters—significantly constrains the solution space. Thus, leveraging single-cell multi-omics data from heterogeneous tissues enables robust estimation of expression metrics across populations, in turn improving parameter inference. This strategy highlights the usefulness of integrating scRNA-seq and scATAC-seq data and allows quantification of regulatory mechanisms at both gene-specific and population-specific levels, offering deeper insights into gene regulation.

## 3 Application to the Dataset PBMC

For our first application of the model, we selected the dataset Peripheral Blood Mononuclear Cells (PBMC) to demonstrate our model s ability to capture gene expression dynamics and regulatory strategies across distinct cell populations.

PBMC is a standard dataset in single-cell studies due to its heterogeneous yet well-characterized cell types. As a paired multi-omics dataset, PBMC provides both gene expression and chromatin accessibility information for each cell. The dataset consists of 10,000 cells grouped into 8 major cell types. To increase the number of distinct populations available for model fitting, we subdivided these cell types into a total of 20 clusters. Each cluster comprises between 100 and 1,000 cells, providing a robust basis for accurately estimating the mRNA expression descriptors and the fraction of open chromatin. Moreover, for this first application of our model, we kept only promoter regions from the ATAC data (see the Discussion for alternative ideas).

Next, we fit the model to the top 500 most expressed genes of the PBMC dataset. We inspected model output by visualizing the comparison between observed and fitted value of *M* ^*e*^ and *F* ^*c*^ for different cell types. s a typical example, we show MAP3K8, a gene that codes for a protein kinase (Figure 4). We observe that our model and inference procedure accurately fit the data and that model captures information across both omics modalities, RNA and ATAC.

**Fig. 4.**
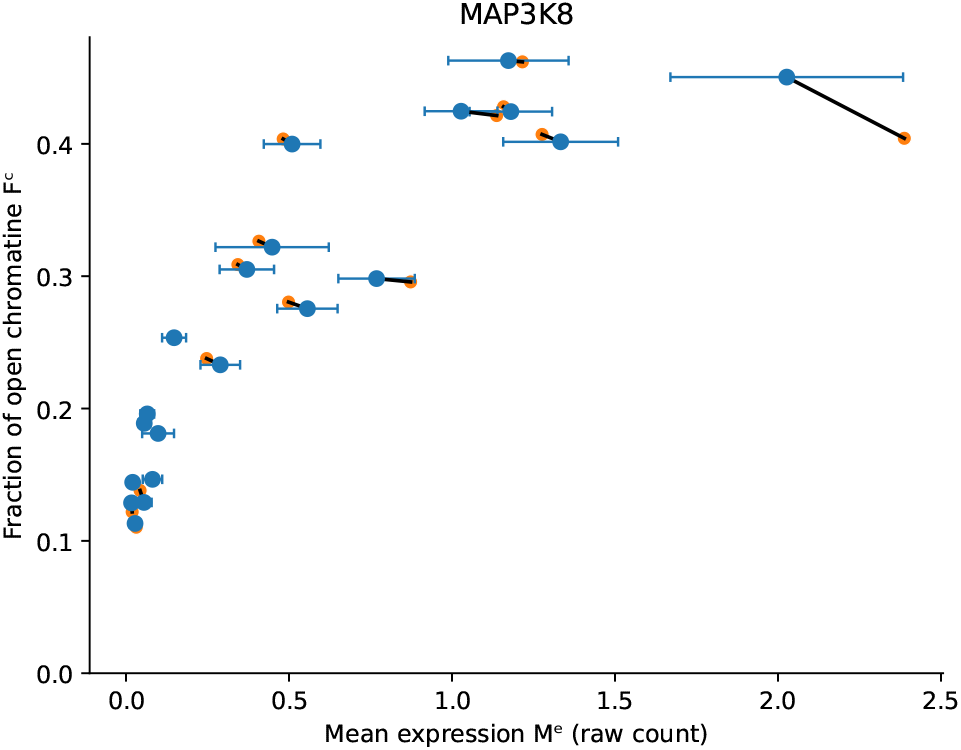
Model performance for the gene *MAP3K8*, comparing *M*^*e*^ and *F*^*c*^ for each cell type. Data is shown as mean expression *±* 95% confidence intervals for *M*^*e*^. Per cell type, model (blue dots with whiskers) and data (orange dots) are linked by a black line to more easily confirm quality of fit.

### 3.2 Interpretation of Model Parameters reveals Expression Strategies

Due to a careful design of the model and the biological hypotheses underlying it, model parameters are readily interpretable. As mentioned above, the model comprises five parameters per gene that are fit to all cell types at once, as well as two parameters that vary by cell type. Thus, for a given gene, there are 5 + 2 × *k* parameters, where *k* is the number of cell types.

The five ‘fixed parameters create a landscape within which the two additional parameters, *λ* (probability to open chromatin) and *β* (probability to have TF bound), are fit to the data per cell type. Generally speaking, a distinct landscape is obtained for each of the model descriptors *F* ^*c*^, *M* ^*e*^ and *V* ^*e*^, where each descriptors imposes its own constraints on a given cell type. At the same time, the optimization process changes the values of the five other parameters to find a landscape allowing a good fit for every pair of *λ* and *β*.

Let us elaborate on the example landscape for *MAP3K8* (Figure 5). For each cell type, the optimization process attempts to find a pair (*λ*^*ct*^, *β*^*ct*^) that fits the value of the three descriptors. Within the landscape of *MAP3K8*, the red lines indicate the regions corresponding to the values of the three CD4+ descriptors. To fit these descriptors well, (*λ*^*CD*4+^, *β*^*CD*4+^) have to be at the intersection of the three red lines. Thus, for a given cell type the possible combinations of (*λ*^*ct*^, *β*^*ct*^) are limited. Also note that as *λ* approaches zero, all three descriptors tend toward zero; if the chromatin never opens, the gene cannot be expressed.

**Fig. 5.**
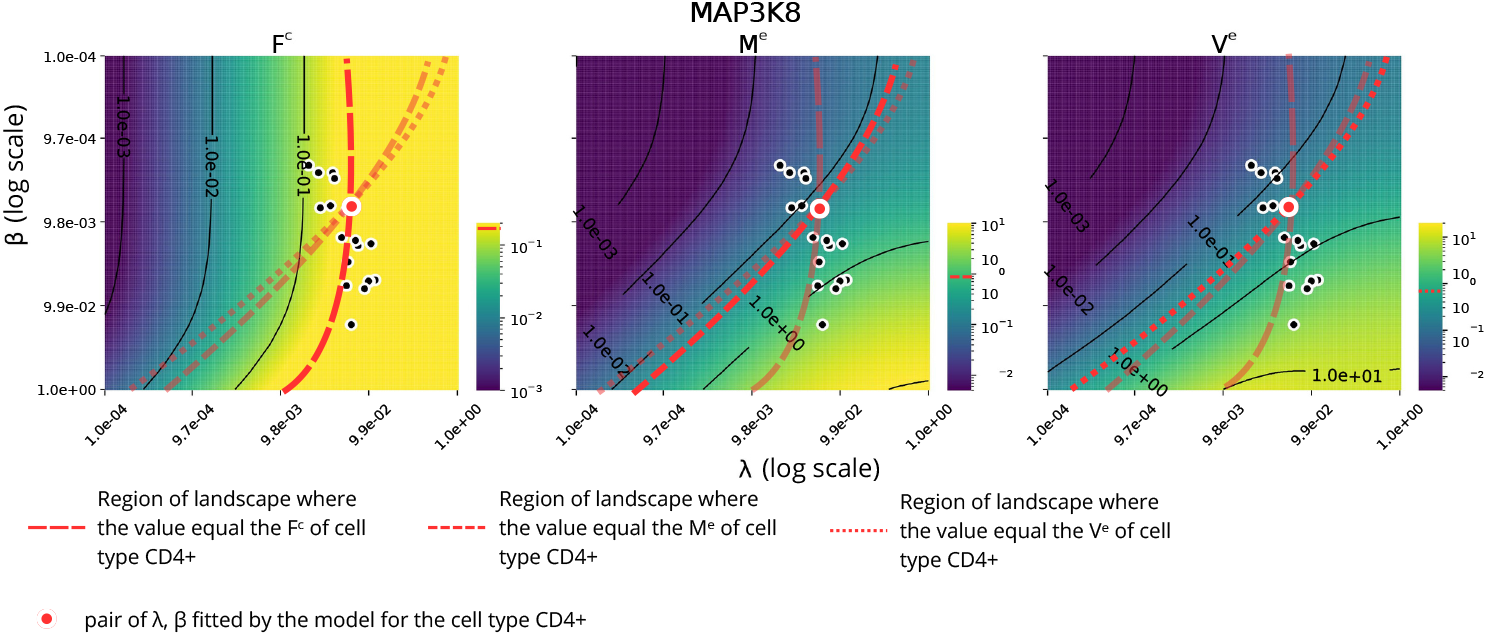
Landscape created by model parameters of the gene *MAP3K8*. Once *µ, υ, ϵ*_*a*_, *ϵ*_*b*_ and *δ* are fitted for a gene, the descriptors *F*^*c*^, *M*^*e*^, and *V* ^*e*^ (left, middle, and right panel) can only vary according to *λ* (probability to open chromatin, x axis) and *β* (probability to have TF bound, y axis). The resulting triptych landscape of a gene gives a context to how cell types are fit (black and red dots). Note the gradient is in logarithmic scale and cell types with *β* = 0 cannot be visualied.

Conversely, when *β* approaches zero, gene expression becomes linear with respect to *λ*. This is the expected behavior of the model as argued in Section 2.1 (see also [18]).

By analyzing the range of parameter values inferred from the model across multiple genes, we can observe distinct means through which gene expression is achieved. We refer to these as Expression Strategies (ES). Once the model is fit to a large set of genes available in the PBMC dataset, we cluster the genes on the basis of their parameter values using a spectral decomposition algorithm [14]. We then interpret the resulting clusters of genes, i.e., parameter profiles, as indicating shared expression strategies.

Each of the seven clusters exhibits a strategy for transcript expression (Figure 6). ES3 is characterized by a low value of *µ*, indicating that genes spend a significant amount of time in the active state. However, this is compensated by a low value of *ϵ*_*a*_, meaning that most transcripts originate from the TF-bound state. ES0, ES2, ES4 and ES5 share a common profile with relatively homogeneous values of *µ, δ, ϵ*_*a*_, and *ϵ*_*b*_, and primarily differ by their *λ* values. ES1 stands out due to its markedly high *δ* value, suggesting a high degradation rate of transcripts. Finally, ES6 is distinguished from the others by a higher *β* value and an elevated basal expression level, reflecting a transcriptional strategy with enhanced chromatin accessibility and constitutive expression. In summary, the seven clusters with each their own expression strategy may be simplified to four distinct ES. In our future work, we will investigate the relationship between these expression strategies and known biological functions or transcription factors.

**Fig. 6.**
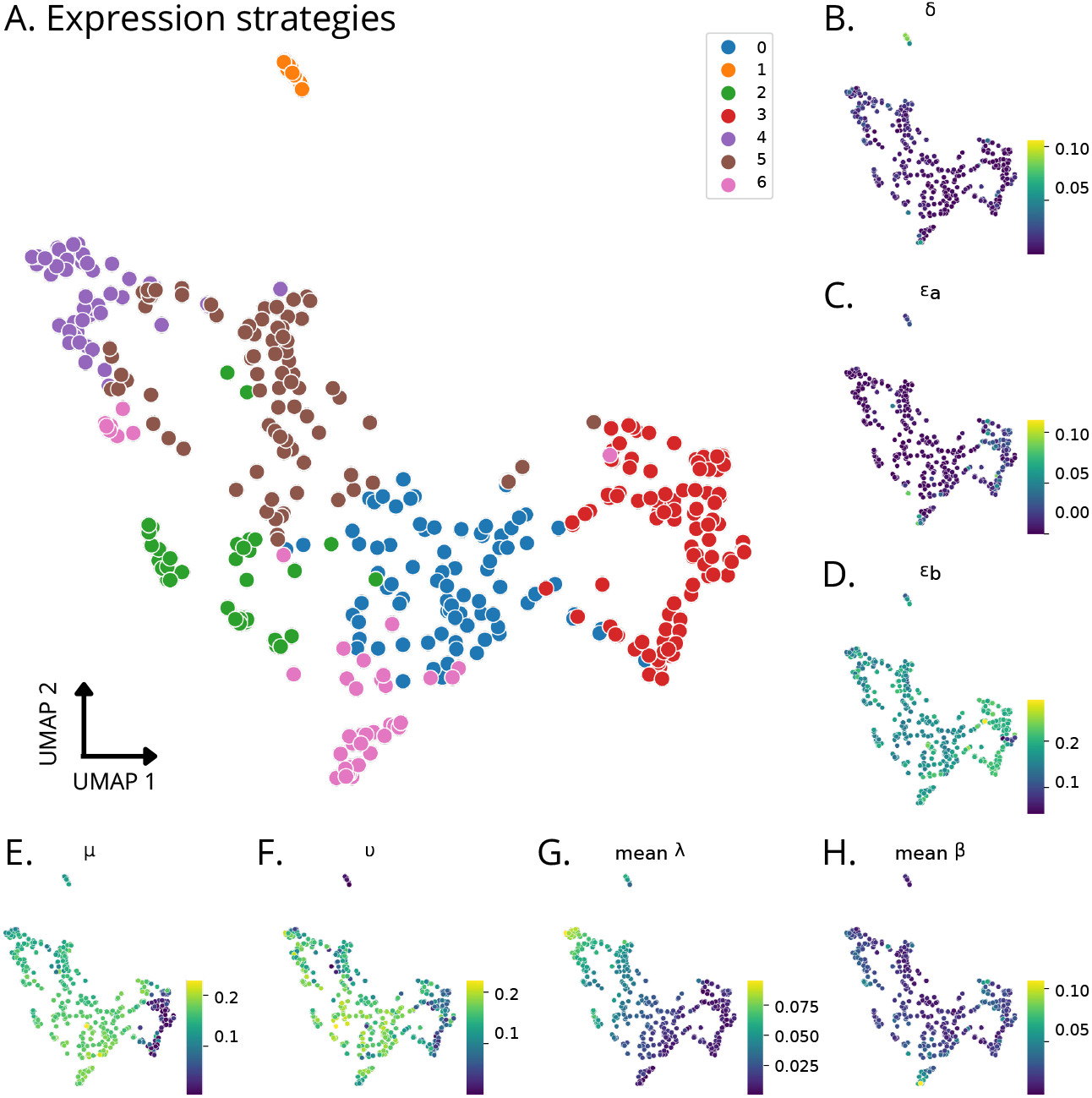
Uniform Manifold Approximation and Projection (UMAP) of genes (dots) on the basis of their model parameters. **A** Seven expression strategies are identified by a spectral decomposition clustering. **B-H** UMAPs colored by model parameter values. For *λ* and *β* mean values over cell types are shown, since these two parameters vary per cell type. The other five parameters do not vary across cell types.

## 4 Discussion

In this work, we introduced a new three-state model for gene expression, that provides a single framework to account for (i) chromatin dynamics via transitions between inactive and active states, and (ii) transcription factor (TF) regulation via transitions between active and TF-bound states [20]. This dual-layer view enables the model to disentangle and quantify the distinct contributions of epigenetic accessibility and TF activity to gene expression.

As a next step, it would be beneficial to revisit the relationship between chromatin dynamics and expression levels [24]. While the current model, inherited from the two-state framework, assumes a linear relationship between burst frequency and expression, preliminary observations suggest that a logarithmic or saturating relationship may better reflect biological reality. Moreover, currently we take into account only chromatin accessibility close to the TSS (promoter) region of a gene. It has been shown to be informative to take into account chromatin accessibility in multiple regions around a TSS [5] and we will explore if such an approach is beneficial for our model.

The use of single-cell multi-omics data, combining RNA-seq and ATAC-seq, offers a powerful strategy to inform the model. These datasets allow the extraction of features such as gene expression mean and variance, as well as chromatin accessibility, across multiple cell populations derived from a single experiment. This improves model fitting and enables gene-wise inference across diverse cellular contexts. However, single-cell data remain prone to technical noise such as dropouts and batch effects [12], which may impact the robustness of the results. Future work should therefore involve larger datasets with increased sample diversity to ensure model generalizability.

The model provides analytical expressions for three key features, i.e., the fraction of open chromatin *F*, and the mean *M* and variance *V* of mRNA transcripts, allowing efficient parameter inference through the proposed loss function. We maintain the number of gene-intrinsic parameters as high as possible to limit the number of feasible model solutions for our data. While this constraint reduces the model s flexibility —potentially impacting the results for certain genes— it enables a more robust interpretation of the results for the remaining genes. Nonetheless, challenges remain in tuning critical hyperparameters, such as the strength of L1 regularization on *β*, which governs TF regulation.

Finally, the interpretability of model parameters allows for a biologically meaningful understanding of the diverse strategies used to regulate gene expression within a given cell type. In the PBMC dataset, clustering based on inferred parameters identified seven groups of genes, revealing four major expression strategies. These strategies differ primarily in their regulation probabilities, burst frequencies, and basal expression levels. However, it remains challenging to ensure that each gene is assigned to a particular strategy due to true underlying biological differences, rather than due to the optimization routine finding a local minimum of the loss function. Addressing this limitation will require further refinement of the inference procedure and potentially the development of additional robustness criteria or model selection strategies to verify the biological relevance of the identified expression strategies.

## Computational Environment

All analyses were performed using Python (version 3.12), with key packages including Scanpy (version 1.9) for single-cell data processing and PyTorch (version 2.3) for model fitting and optimiation. Computations were carried out on a Dell laptop equipped with an Intel Core i9-13900H 13th generation CPU, a NVIDIA RTX A1000 GPU, and 32 GB of RAM.

## Dataset

The PBMC single-cell multi-omics dataset was obtained from 10x Genomics (PBMCs from C57BL/6 mice (v1, 150×150), Single Cell Immune Profiling Dataset by Cell Ranger v3.1.0, 10x Genomics, (2019, July 24)”).

## Code Availability

The code used for the model implementation and training procedures is publicly available at: https://github.com/PeyricT/3StatesGeneExprModel.

## Acknowledgments

This study was funded by PEPR Santé Numérique, project AI4scMed, France 2030 ANR-22-PESN-0002.

## A Analytical solution for Fractions, Means and Variances

To effectively reduce the computational cost of fitting the model, we derive the mean and variance of gene expression as well as the fraction of open chromatin as analytical functions of model parameters, rather than relying on numerical simulations. To achieve this, we need to compute the first and second moments of the moment-generating function derived from the distribution predicted by the model. This process begins by solving the Kolmogorov system obtained from the continuous-time formulation of the model. We then make use of three so-called generating functions. Subsequently, we reframe the problem in a vectorial perspective, which enables the efficient computation of the exponential moments of the resulting function.

### A.1 Solving Markov Process in Continuous Time at Equilibrium

The transitions between the state space of the three-state model define a birth and death process. We have three Markovian processes in continuous time, one for each Markov state. Let *p*_*i,n*_(*t*) be the probability to be in state *i* with *n* transcripts at time *t*. According to our model, we have (note that this is not at equilibrium and with the convention *p*_*i*,−1_ = 0):

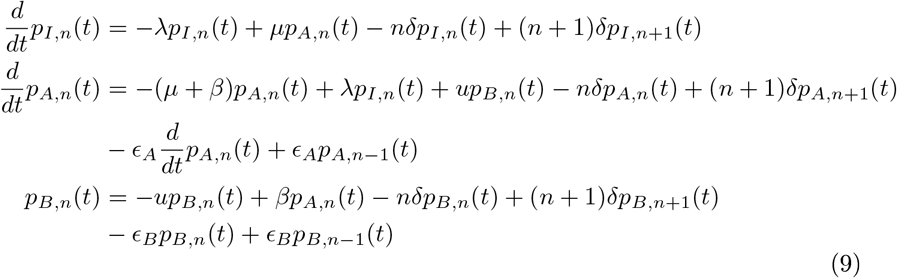

The observable output (i.e., the moments of the transcript-count distribution we are computing) of the model is given by the following equation:

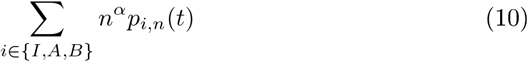

with *α* = 1 for the mean number of transcript at time t,

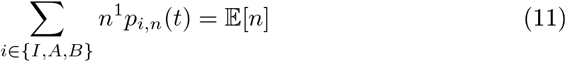

and *α* = 2 the second raw moment, from which one can get the variance via

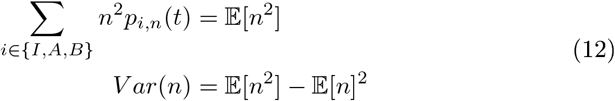

In similar fashion as demonstrated by Peccoud *et al*. [17], we solve the Kolmogorov system associated with this process. Which will allow us to reformulate and analyze the following system of differential equations:

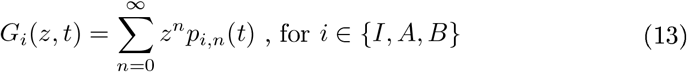

Thus our observable is now

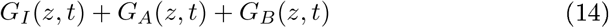

We can finally arrive at the following system satisfied by the generating functions (13) on the system (9):

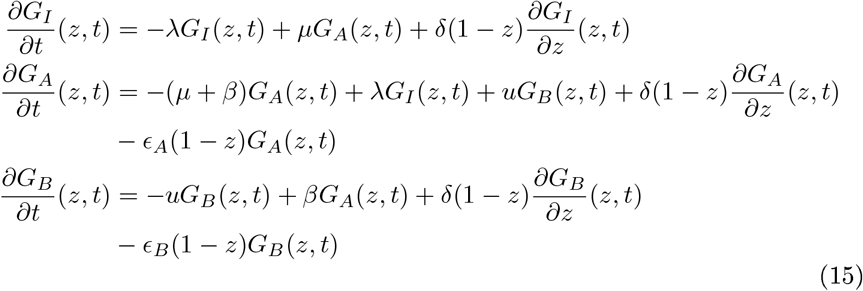

We can now proceed to solve this system at equilibrium. To do so, we adopt the vectorial point of view.

### A.2 Re-framing the System as a Vectorial Expression

Here we compute the mean and variance of the distribution generated by the model at equilibrium (canceling the time derivatives in (9) and (13)). To achieve this, we focus on deriving the generating function corresponding to equilibrium. Through the Kolmogorov equation, it is easy to see that we have a simple system on the quantities *G*_*i*_(1, *t*) = ∑ _*n*_ *p*_*i,n*_(*t*)

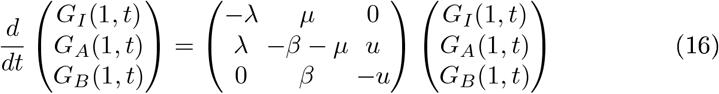

For equilibrium, since we have non-negativity, the values are give by the stationary distribution of the corresponding simple Markov process. This corresponds to the eigenvector of the transition matrix associated to eigenvalue 0. We can solve easily

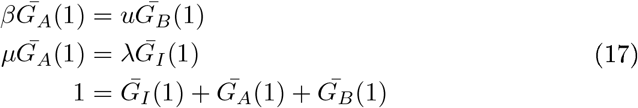

which results in:

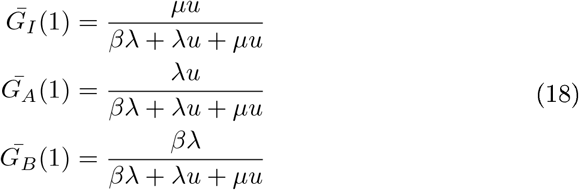

Expanding the moment generating function in terms of powers of *z* − 1 rather than *z* leads to (Taylor expansion)

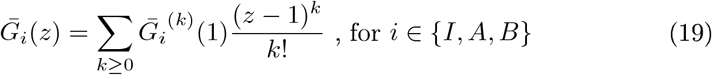

Now we can see the previous system as a vectorial space with:

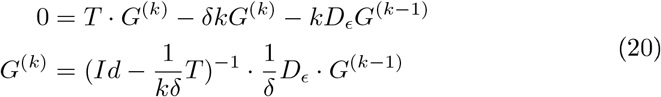

with transition matrix, (as introduced above)

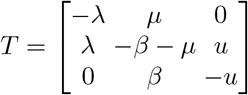

and expression matrix

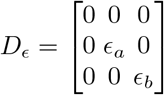

It follows that:

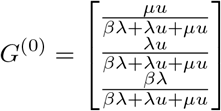

Finally, *F* ^*c*^ is defined as the fraction of time that chromatin is open, in other words, the probability to be in state A or B. This means the analytical solution is:

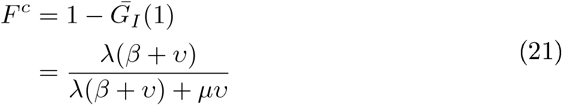

### A.3 Mean and Variance Equation using Exponential Moments

A small difficulty is to switch between expansion in terms of powers of *z* and *z* − 1. By using a power series, we can easily compute the exponential moments [17] corresponding to the evaluation of the *k*−th derivative at *z* = 1.

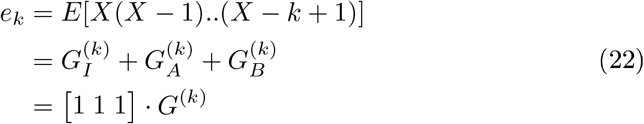

Thus, the mean of the asymptotic distribution is

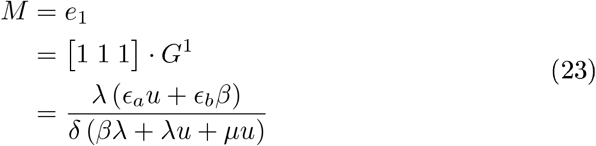

which we will use as the mean mRNA expression (*M* ^*e*^).

The mathematical expression for the variance is more complex and lengthy, so we introduce two auxiliary variables, *a* and *b*, to simplify the equation:

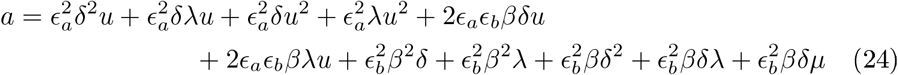

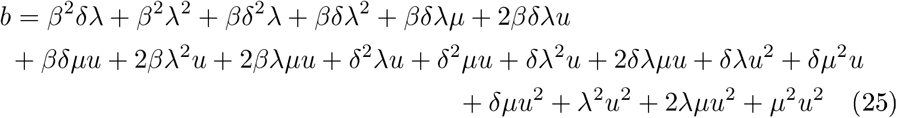

Using *a* and *b*, we can write:

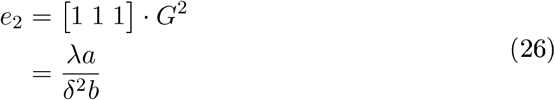

Thus, the variance of the asymptotic distribution is

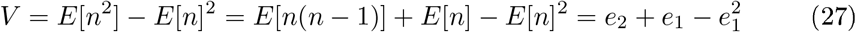

We use this variance as the mRNA expression variance (*V* ^*e*^). This completes our analytical computation.

